# Assessing the motivational effects of ethanol in mice using a discrete-trial current-intensity intracranial self-stimulation procedure

**DOI:** 10.1101/731612

**Authors:** Amanda M. Barkley-Levenson, Andre Der-Avakian, Abraham A. Palmer

## Abstract

**Background:** Alcohol (ethanol) produces both rewarding and aversive effects, and sensitivity to these effects is associated with risk for an alcohol use disorder (**AUD**). Measurement of these motivational effects in animal models is an important but challenging aspect of preclinical research into the neurobiology of AUD. Here, we evaluated whether a discrete-trial current-intensity intracranial self-stimulation (**ICSS**) procedure can be used to assess both reward-enhancing and aversive responses to ethanol in mice.

**Methods:** C57BL/6J mice were surgically implanted with bipolar stimulating electrodes targeting the medial forebrain bundle and trained on a discrete-trial current-intensity ICSS procedure. Mice were tested for changes in response thresholds after various doses of ethanol (0.5 g/kg-1.75 g/kg), using a Latin square design.

**Results:** A 1 g/kg dose of ethanol produced a significant reward-enhancement (i.e., lowered response thresholds), whereas a 1.75 g/kg dose produced an aversive effect (elevated response thresholds).

**Conclusions:** The discrete-trial current-intensity ICSS procedure is an effective assay for measuring both reward-enhancing responses to ethanol as well as aversive responses in the same animal. This should prove to be a useful tool for assessing the effects of experimental manipulations on the motivational effects of ethanol in mice.

---

Alcohol (ethanol) is a compound stimulus with both rewarding and aversive effects (Verendeev and Riley 2013), and motivation to consume ethanol is influenced by sensitivity to these effects. In humans, the rewarding effects of ethanol include euphoria, stimulation, and drug-liking; in contrast, aversive effects include sedation, dysphoria, and drug-disliking (Rueger and King 2013; Fridberg et al. 2017). Both greater sensitivity to ethanol’s rewarding effects and reduced sensitivity to ethanol’s aversive effects have been associated with a greater risk of an alcohol use disorder (AUD) (e.g. Schuckit et al. 2007, King et al. 2014). Understanding the neurobiological basis of ethanol’s motivational effects may therefore provide critical insight into excessive ethanol consumption and AUD. Furthermore, drugs that are intended to treat AUD could potentially act via blocking the rewarding effects of ethanol or by enhancing the aversive effects of ethanol.

The intracranial self-stimulation (ICSS; sometimes called brain reward stimulation) procedure provides a direct measure of brain reward function, where responding is reinforced by electrical stimulation of the mesolimbic reward circuitry. Response thresholds can be established by determining the level of stimulation that maintains operant responding. Drugs of abuse can enhance the rewarding properties of this stimulation, presumably through dopaminergic activation (Carlezon and Chartoff 2007). This reward-enhancement effect is characterized by a lowering of ICSS response thresholds, wherein less stimulation is required to maintain responding. In contrast, increases in ICSS response thresholds are interpreted as an aversive or anhedonic response because a greater stimulation intensity is required to maintain the same level of responding. ICSS has been widely used to study the reward-enhancing properties of drugs of abuse, with nearly all classes of abused drugs showing enhancement of reward in ICSS procedures (for review, see Kenny et al. 2018; Negus and Miller 2014). However, ICSS has not been routinely used with ethanol in rodents, and especially has been underutilized in mice. To the best of our knowledge, the only previous efforts in mice have also exclusively used rate-frequency procedures (Fish et al. 2010, 2012) that measure drug-induced increases in response rate. These procedures can be more sensitive to locomotor changes (both stimulation and sedation), and therefore may present challenges for interpreting results across ethanol doses. Here, we demonstrate that a discrete-trial current-intensity procedure to measure ICSS response threshold current can be used to assess both reward-enhancing and aversive responses to ethanol in the same animal across a range of doses.

## Methods

### Animals and husbandry

Mice were tested in two cohorts approximately six months apart. Adult male and female C57BL/6J mice were bred in house (cohort 1) or purchased from Jackson Laboratories (cohort 2). Mice were housed 2-5 per cage and had *ad libitum* access to food (Envigo 8604, Indianapolis, IN) and water. Mice were housed on a 12h:12h reverse light-dark cycle with lights off at 07:00h, and all testing occurred during the dark phase. All procedures were conducted at the University of California San Diego (**UCSD**) and were approved by the UCSD Institutional Animal Care and Use Committees and were conducted in accordance with the NIH Guidelines for the Care and Use of Laboratory Animals.

### Surgical procedures

ICSS surgical procedures have been described previously (Stoker et al. 2008). Briefly, mice were surgically implanted with a bipolar stimulating electrode (Plastics One Inc., Roanoke, VA) aimed at the medial forebrain bundle (A/P: −1.58, M/L: ±1.0, D/V: −5.3; Paxinos and Franklin 2001). Left/right position of the electrode was counterbalanced across animals. Electrodes were secured to the skull with dental acrylic and two stainless steel anchor screws. All animals were allowed to recover for one week prior to the start of behavioral training.

### ICSS apparatus

ICSS training and testing were conducted in eight Plexiglas operant chambers (30.5 × 24 × 27 cm; Med Associates, St. Albans, VT). Each operant chamber was enclosed within a light and sound-attenuated chamber (40 × 60 × 63.5 cm). Intracranial stimulation was delivered by constant current stimulators (Stimtek model 1200c, San Diego Instruments, San Diego, CA). The mice were connected to the stimulation circuit through flexible bipolar leads (Plastics One, Roanoke, VA) attached to gold-contact swivel commutators (model SL2C, Plastics One, Roanoke, VA) mounted above the operant chamber. The operant response required by the subjects was a quarter turn of a wheel manipulandum (5.5 cm diameter, 4 cm width) that extended 1.5 cm from one wall of the operant chamber.

### ICSS procedure

The ICSS procedure was adapted from a discrete-trial current-intensity threshold procedure used in rats (Kornetsky and Esposito 1979; Markou and Koob 1992) and has been described previously for mice in conjunction with other drugs (Stoker et al. 2008; Stoker and Markou 2011; Gill et al. 2004). Briefly, mice were first trained to turn a wheel manipulandum on a fixed-ratio 1 (**FR1**) schedule of reinforcement to receive electrical stimulation. The frequency of the electrical stimulation was fixed (100 hz), and the current was adjusted for each animal to maintain responding (120-180 μA). After acquisition of the FR1 schedule (i.e., two consecutive test sessions with 200 reinforcers earned in less than 10 min), animals were trained on the discrete-trial current threshold procedure. Each trial was initiated by the delivery of a noncontingent electrical stimulation followed by a 7.6 s response window within which the subject could make a response to receive a second contingent stimulation. The electrical current was varied across descending and ascending series of trials to determine the minimal current level that maintained responding (i.e., response threshold). Mean response latency (time between trial start and response for the contingent stimulation) was also recorded.

### Experimental design

Following training, mice received one test session per day. Mice were given at least 10 sessions to establish stable baseline response thresholds before the start of ethanol testing. An animal was considered to have a stable initial baseline when there was less than 10% variation in thresholds over five consecutive sessions. Mice were tested on a series of ethanol doses using a Latin square design. On ethanol test days, mice were weighed and given an i.p. injection of the assigned ethanol dose. They were returned to the home cage for 5 min and then tested in the discrete-trial current-intensity ICSS procedure, which lasted approximately 20-35 min. Stable baseline thresholds were reestablished between each ethanol test session (less than 10% variation in threshold over at least three consecutive sessions).

### Drugs

Ethanol solution (Deacon Laboratories Inc., King of Prussia, PA) was made fresh each test day (20% v/v in saline) and administered i.p. at doses of 0 g/kg [saline], 0.5 g/kg, 0.75 g/kg, 1 g/kg, 1.5 g/kg, and 1.75 g/kg. This dose range was based on the effective dose range seen in previous studies (Fish et al. 2010) and pilot experiments in our laboratory that indicated higher doses produced significant locomotor sedation that interfered with task performance.

### Analyses

All analyses were conducted using SPSS (version 25; IBM, Armonk, NY). Response threshold on each ethanol test day was converted to a percent of baseline threshold for each individual animal. Because not all animals completed each stage of the Latin square due to technical issues (e.g. loss of head cap, electrode chewed by cagemates), repeated measures analysis could not be reliably conducted; instead we analyzed the data by ANOVA with ethanol dose, sex, and cohort as factors for both response threshold and mean response latency. Group sizes were n=5 for 1.5 g/kg, n=7 for 1 g/kg, and n=6 for all for all other doses. Because we were specifically interested in which ethanol doses would produce a significant change compared to control (saline), we also used planned comparisons (two-sample t-tests) of each ethanol dose vs. saline. The level of significance was 0.05.

## Results

Figure 1 shows the ethanol dose-response curve for ICSS response thresholds. Analysis of response thresholds showed a main effect of ethanol dose (*F*_5,18_=4.74, p<0.01). There were no main effects of sex or cohort, and no significant interactions. Planned comparisons of response threshold for each ethanol dose to saline showed that the 1 g/kg dose significantly reduced thresholds compared to saline (two-tailed: *t*_11_=2.27, p<0.05). This is consistent with a reward-enhancing effect. In contrast, the 1.75 g/kg dose significantly increased thresholds compared to baseline (two-tailed: *t*_10_=-2.48, p<0.05), which is consistent with an aversive effect. No other dose groups were found to differ significantly from saline (p>0.1 for all). The threshold for one mouse at the 1 g/kg dose was found to be a statistical outlier (>2 standard deviations higher than the mean) and this animal’s 1g/kg data were excluded from analyses.

**Figure 1.**
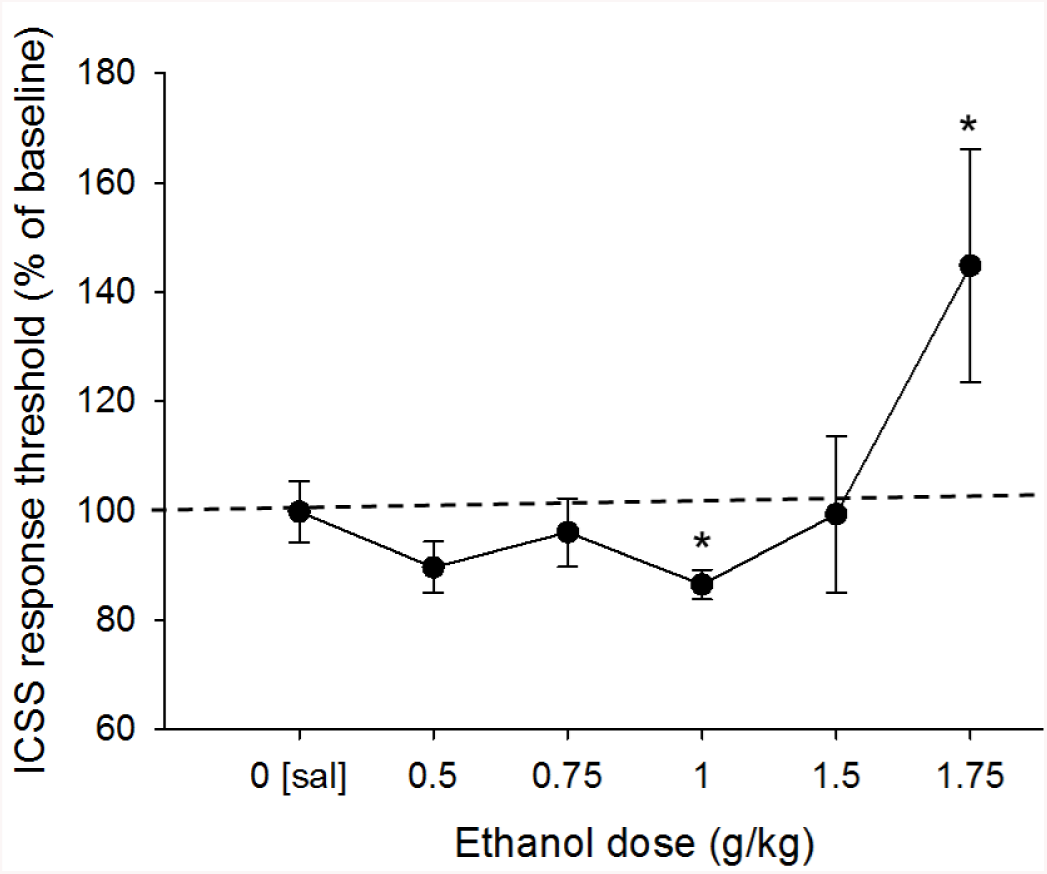
ICSS response thresholds across ethanol dose. Thresholds are presented as a percent of baseline (dashed line). * indicates statistically significant difference from saline group (p<0.05). N=5-7 per dose.

Figure 2 shows response latencies at each dose. There was a main effect of dose on response latencies (*F*_5,18_=9.53, p<0.001). Planned comparisons showed that the 1 g/kg (*t*_11_=-2.74, p<0.05), 1.5 g/kg (*t*_9_=-2.94, p<0.05), and 1.75 g/kg (*t*_10_=-3.81, p<0.01) doses of ethanol significantly increased latencies compared to saline. No other doses significantly differed from saline (p>0.1 for all), and there were no other main effects or significant interactions.

**Figure 2.**
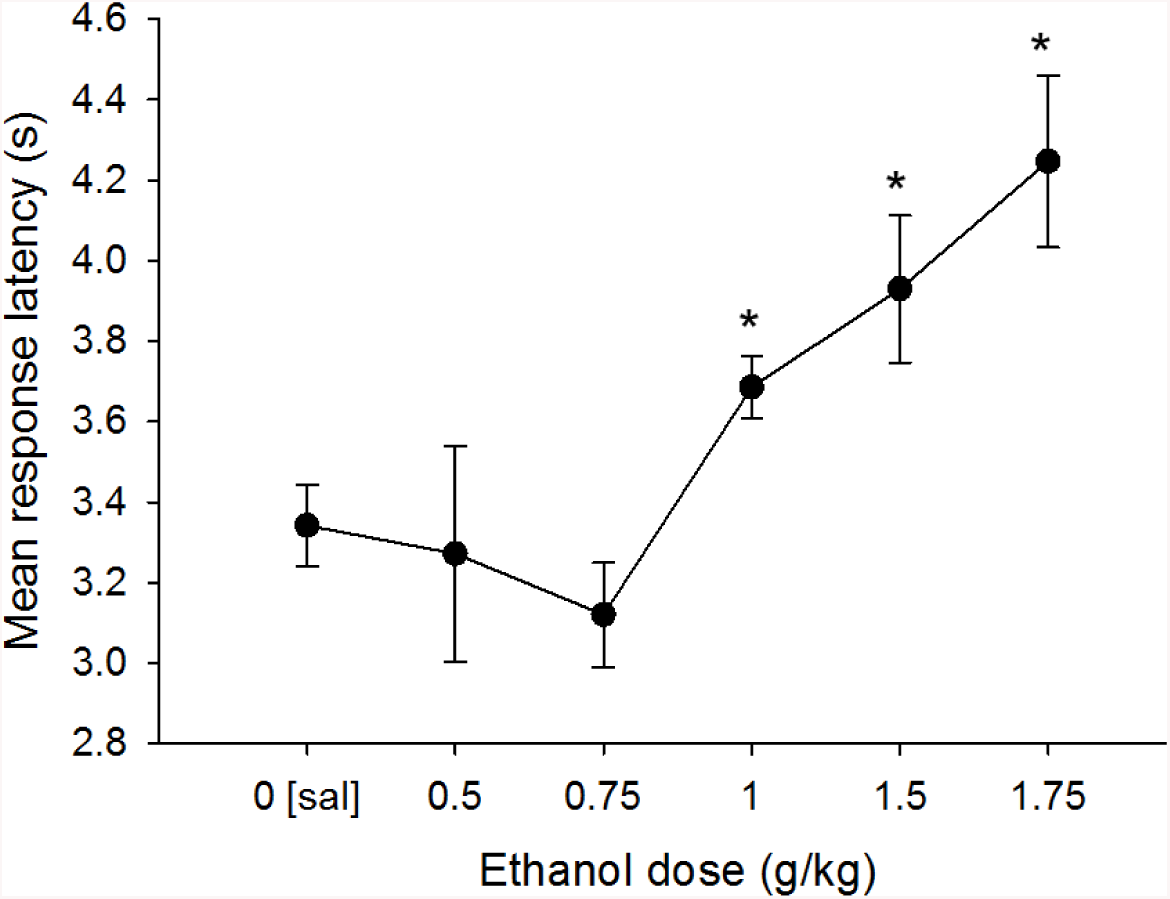
Mean response latencies across ethanol doses. Mean response latencies across all trials in a session are shown for each dose of ethanol. * indicates significant difference from saline group. N=5-7 per dose.

## Discussion

This experiment provides the first data showing that a discrete-trial current-intensity ICSS procedure can assess both reward-enhancing and aversive effects of ethanol in mice. We demonstrated that a 1 g/kg dose of ethanol produces a modest but significant decrease in response thresholds, indicating a reward-enhancing effect. Treatment with a 1.75 g/kg dose produced a significant increase in response thresholds, which is consistent with an aversive response. We observed longer response latencies at the 1 g/kg, 1.5 g/kg, and 1.75 g/kg doses, which may indicate locomotor sedation or the presence of competing behaviors. However, all mice were able to successfully complete the session at these doses and it is therefore unlikely that locomotor deficits account for the observed changes in thresholds.

We are only aware of two prior mouse studies that involved ethanol and ICSS. One previous study using a rate-frequency procedure in C57BL/6J and DBA/2J mice found that a 0.6 g/kg dose had a reward-enhancing effect at 0-15 min post-treatment in C57BL/6J mice but had no effect at later time points (Fish et al. 2010). Doses from 0.6 g/kg-1.7 g/kg were found to be reward-enhancing in DBA/2J mice for up to 30 min post-treatment., but also produced significant increases in maximum response rate. In a different study using mice that had been selectively bred for high (FAST) or low (SLOW) locomotor response to ethanol, doses up to 2.4 g/kg were found to be reward-enhancing for the FAST mice, whereas there were no doses that produced significant threshold changes in the SLOW mice (Fish et al. 2012). The effective dose range for assessing reward-enhancing vs. aversive effects is likely genotype-specific and therefore the doses identified in our study should be re-established when working with non-C57BL/6J mice.

There are also several procedural variations between our study and the few previous published reports on ethanol ICSS in mice that should be noted. In our experiment, we used an i.p. injection as the route of administration, whereas others have used intragastric administration via oral gavage (Fish et al. 2010, 2012). This could potentially explain some of the differences we noted in the dose-response and time course, although one previous study in rats found no difference between i.p. and intragastric administration, with both routes of administration producing no effect on thresholds (Schaefer and Michael 1987). Our study also used the discrete-trial current-intensity procedure instead of a rate-frequency procedure. Both procedures are intended to assess the same underlying process, but the discrete-trial current-intensity procedure may be more resistant to drug-induced changes in rate of responding and nonspecific performance deficits (Markou and Koob 1992). In the present experiment we also included both male and female mice; while no sex effects or interactions were observed, our study was likely underpowered to detect such effects.

Previous work with rats has shown mixed effects of ethanol on ICSS, with some studies showing decreases in thresholds (Lewis and June 1994; Moolten and Kornetsky 1990) and some studies showing no change or increases in thresholds (Carlson and Lydic 1976, Schaefer and Michael 1987). Previous mouse studies have shown decreases in thresholds or no effect depending on the genotype tested, although some doses do appear to produce non-statistically significant increases in thresholds (Fish et al. 2010, 2012). Our findings appear to be unique in identifying different doses that can either decrease (1.0 g/kg) or increase (1.75 g/kg) response thresholds. This suggests that our procedure may be useful for evaluating both reward-enhancing and aversive sensitivity to ethanol in the same animal. It should also be noted that the time course of ethanol effects on ICSS response threshold may prove to be an important variable. In the present study, response thresholds were averaged over the entire session which covered roughly 25-40 min. However, it is possible that the strength and/or direction of ethanol’s effects on response thresholds may vary within this time period. Future experiments will be needed to determine whether this is the ideal time window for observing these effects.

In summary, we have presented the first evidence that ethanol modulates response thresholds in a discrete-trial current-intensity ICSS procedure in a dose-dependent manner in mice. This procedure has the potential for broad application in ethanol research in order to better understand the neurobiology of ethanol’s motivational effects.

## Contributors

All authors contributed to the design of this study. AMB-L was responsible for data collection and analysis. AMB-L drafted the manuscript with input from AD-A and AAP. All authors approved the final version of the manuscript for submission.

